# Lateralized Sensorimotor Evoked Potentials during Visuomotor Transformation in Real and Imagined Movements

**DOI:** 10.1101/2023.05.24.542085

**Authors:** Nikolay Syrov, Lev Yakovlev, Alexander Kaplan, Mikhail Lebedev

## Abstract

The neural mechanisms underlying motor preparation have attracted much attention, particularly because of the assertion that they are similar to the mechanisms of motor imagery (MI), a technique widely used in motor rehabilitation and brain-computer interfaces (BCIs). Here we clarified the process of visuomotor transformation for the real and imagined movements by analyzing EEG responses that were time locked to the appearance of visual targets and movement onsets. The experimental task required responding to target stimuli with button presses or imagined button presses while ignoring distractors. We examined how different components of movement-related potentials (MRPs) varied depending on the reaction time (RT) and interpreted the findings in terms of the motor noise accumulation hypothesis. Furthermore, we compared MRPs and event-related desynchronization (ERD) for overt motor actions versus motor imagery. For the MRPs, we distinguished lateralized readiness potentials (LRPs) and reafferent potentials (RAPs). While MRPs were similar for the real and imagined movements, imagery-related potentials were not lateralized. The amplitude of the late potentials that developed during motor imagery at the same time RAPs occurred during real movements was correlated with the amplitude of β-ERD. As such they could have represented sensorimotor activation triggered by the imagery. LRPs that occurred during real movements lasted longer for longer RTs, which is consistent with activity accumulation in the motor cortex prior to overt action onset. LRPs occurred for non-target stimuli, as well, but they were small and short lived. We interpret these results in terms of a visuomotor transformation, where information flows from visual to motor areas and results in a movement, a decision not to move and/or a mental image of a movement. The amplitude of the late positive peak that developed during MI was correlated with the amplitude of the β-ERD. Since the latency of this component was consistent with the timing of RAP, we suggest that it is a non-lateralized RAP-like component associated with sensorimotor activation during kinesthetic MI.

## INTRODUCTION

Voluntary movements have a complex neuronal mechanism whose dynamics could be described as a sequence of stages, including motor preparation, execution and feedback processing (Bernstein, 1947; Rahman et al., 2019).

Motor preparation is one of the most mysterious stages because of the diversity of vaguely defined covert factors that could motivate and trigger movements (Jennings and van der Molen, 2005). EEG-derived event-related potentials (ERPs) are widely used in sensory-motor tasks, because they provide tracing of the neural processing time course with high temporal resolution (Vaughan, 1969; Berchicci et al., 2016). Movement-related potentials (MRPs) that are time-locked to sensory cues and/or movement onsets are indicative of the sensorimotor integration stages within the action preparation and performance.

The consecutive MRP stages are typically described as a slow readiness potential (RP) followed by a lateralized readiness potential (LRP) and a post-movement reafferent potential (RAP). The first two potentials are related to preparatory processes. The readiness potential (RP), or Bereitschaftspotential (Neafsey, 2021), is not specific to motor behavior per se. Libet (Libet, 1985) observed RPs develop in human subjects many hundreds of milliseconds prior to movement onset and even awareness to act and interpreted these cortical potentials as correlates of unconscious action preparation. It has been also suggested that RPs could represent anticipation without specific encoding of motor parameters or even artifacts related to averaging of noise (Neafsey, 2021). By contrast to RP, LRP is action specific. It develops ∼200ms prior to a movement as a slight contralateral negativity, which corresponds to the timing of conscious decision to move (Trevena and Miller, 2002; Neafsey, 2021).

The mechanisms and functions of LRP are still not well understood. Visual triggers of movements are typically used in the studies of this component (Hackley and Valle-Inclan, 1998; Hohlefeld et al., 2011) but it is usually interpreted in terms of motor processing rather than a visual response (Misirlisoy and Haggard, 2014; Schmitz et al., 2019). In terms of stimulus-response mapping, LRP is an intermediate component in the process of stimulus transformation into a motor response, where a stimulus drives accumulation of noisy activity until a decision threshold is reached (Van Vugt et al., 2014). LRP is affected in many sensorimotor disorders (Bai et al., 2006; Luck et al., 2009; Saville et al., 2015). Thus, the discovery of the nature of this EEG marker of action preparation is an important challenge to be accepted.

Another MRP also known as RAP, occurs after movement onset as a positive contralateral peak (Vaughan, 1969). Because RAP occurs when the movement is already ongoing (and as follows from the name), movement-related afferent signals (tactile and kinesthetic) significantly contribute to these cortical potentials (Bötzel et al., 1997; Berchicci et al., 2016). RAP could also have a more complex role, namely action monitoring, which is different from merely processing afferent inputs. During such monitoring, afferent signals are compared with their mental expectation formed in advance of movement onset (Bernstein, 1947; Greenwald, 1970). RAPs could have a role in this comparison (Verleger et al., 2006; Verleger et al., 2014).

In addition to ERPs, event-related desynchronization (ERD) is a neural correlate of sensorimotor processing that also has a wide practical application in BCIs. EEG spectral power is reduced in the α- and β-frequency bands during the activation of somatosensory and motor cortices (Graimann et al., 2002; Neuper et al., 2006). This ERD occurs during action preparation (2-3 sec before the action onset), as well as during action execution (Pfurtscheller et al., 2002). The relationship between ERD and ERPs remains a subject of discussion (Fairhall et al., 2007; Rogge et al., 2022). For example, anticipatory ERD could develop earlier than LRP and could be even observed in the absence of LRP (Bai et al., 2006; Rogge et al., 2022). In addition, ERD can occur in the absence of overt motor behavior, such as during motor imagery (MI), when movement-related kinesthetic sensations are generated mentally without actual muscle activation (Pfurtscheller, 2000). MI is somewhat similar to motor preparation, with a difference that a movement never starts (Jeannerod, 2001; Jeannerod, 2006). With this respect, Marc Jeannerod proposed the motor simulation theory (MST) where he suggested that MI shares the neuronal substrates with actual movement (Jeannerod 1994; Jeanerood 2001; Glover and Dixon, 2013). This suggestion was supported by many neuroimaging and neurostimulation studies showing that the same neural structures are activated during MI and the actual movements (Schnitzler et al., 1997; Fadiga et al., 1998; Grosprêtre et al., 2016; Mehler et al., 2019). Yet, the mechanisms of MI are not well understood and regardless of its wide acceptance of MST, it is not devoid of criticism (O’Shea and Moran, 2017). Several studies point to a number of differences in how cortex is activated during MI and versus overt motor behavior (Rodriguez et al., 2009; Gabbard et al., 2009; Glover et al., 2020; Caldara et al., 2004; Yashin et al., 2023). ERP studies of MI have yielded inconsistent results. Galdo-Alvarez et al. (2016) reported reduced lateralized readiness potentials during motor imagery compared to actual execution, and Hohlefeld et al. (2011) did not observe any motor MI-related lateralized components.

At the same time, several studies report strong LRPs during MI of both upper and lower extremities (Sosnowska et al., 2021; Carrillo-de-la-Pena et al., 2006). Comparing MI-related ERP components with MRPs enhances our understanding of the subtle distinctions between imagining a movement, planning and executing a movement, and planning a movement but choosing to halt the intended action. In the present study, we investigated the aforementioned EEG patterns under the conditions typically used in P300 BCI tasks, where a participant mentally responds to target stimuli while disregarding distractors (Wolpaw et al., 2022). Two buttons served as potential targets, and participants pressed the target button or imagined this action in response after the button was highlighted; the non-target stimuli had to be ignored. For these conditions, we analyzed target and non-target cortical potentials developing during MI and motor execution (ME) tasks. Furthermore, for the ME task, we analyzed the relationship between LRP characteristics and the parameters of motor performance that could reflect LRP accumulation timing (Van Vugt et al., 2014) such as reaction time. Next, we explored post-movement EEG components, namely RAP, and β-rhythm desynchronization. Since the pre-movement α/β-ERD and MRPs were already matched in many studies, RAP and sensorimotor rhythmic power were not directly matched in any study (for both real imagined movements). Our expectation to find EEG modulations during both MI and ME was partially fulfilled, except for the lack of lateralization in MI-related ERPs. As we found significant β-ERD in MI trials, the value of which correlated with the RAP-like potential, we propose that MI-related ERPs, as they reflect sensorimotor activation, may be useful for the development of BCIs for motor rehabilitation. In addition, we discuss the results obtained in motor execution trials in the context of theories that provide explanations for the neural mechanisms of stimulus-response mapping and motor control.

## METHODS

### Participants

Seventeen healthy volunteers (mean age 23 years, SD=4, 6 female) participated in the study. All were informed of their rights and gave informed consent to participate in the study. The experimental procedures were approved by the Bioethics Committee of the Lomonosov Moscow State University (protocol no. 111-ch). The study followed the Declaration of Helsinki ethical principles for medical research involving human subjects. All subjects were informed of the study procedures and gave written consent to participate.

### Experimental design

The subjects were seated at a table in a comfortable chair with their hands resting on a panel with two buttons (5 cm in diameter and 10 cm in height). The right and left buttons were dedicated for the right and left hand actions, respectively. During the experimental sessions, the buttons were highlighted using built-in LEDs in a semi-random sequence. The participants were instructed to pay attention only to the highlights of the button designated as the target and press that button as quickly as possible in the ME task or imagine performing a button press in the MI task. Mental counting (MC) of the target stimuli served as a control condition. A block design was used, with each task being performed in a separate session (see Figure 1). The order in which these sessions occurred was randomized across the participants. Each session consisted of 6 consecutive runs. Each run started with the presentation of the instruction word "right" or "left" pointing to the target button. The instructions were presented for 5 seconds on a 22-inch LCD monitor placed in front of the subject. Highlights of the target button required an action (ME, MI or MC) to be performed, whereas non-target stimuli had to be ignored. The target button was randomly selected for each run. The stimulus sequence within each run consisted of 30 trials, each of 1000 ms long: 200-ms button highlight followed by 800-ms interstimulus interval. Target and non-target were equal in numbers and randomly intermixed. These settings resulted in 90 target (45 right + 45 left) and 90 non-target responses collected for each task in a given participant. The experimental design is graphically illustrated in Fig.1.

**Figure 1.**
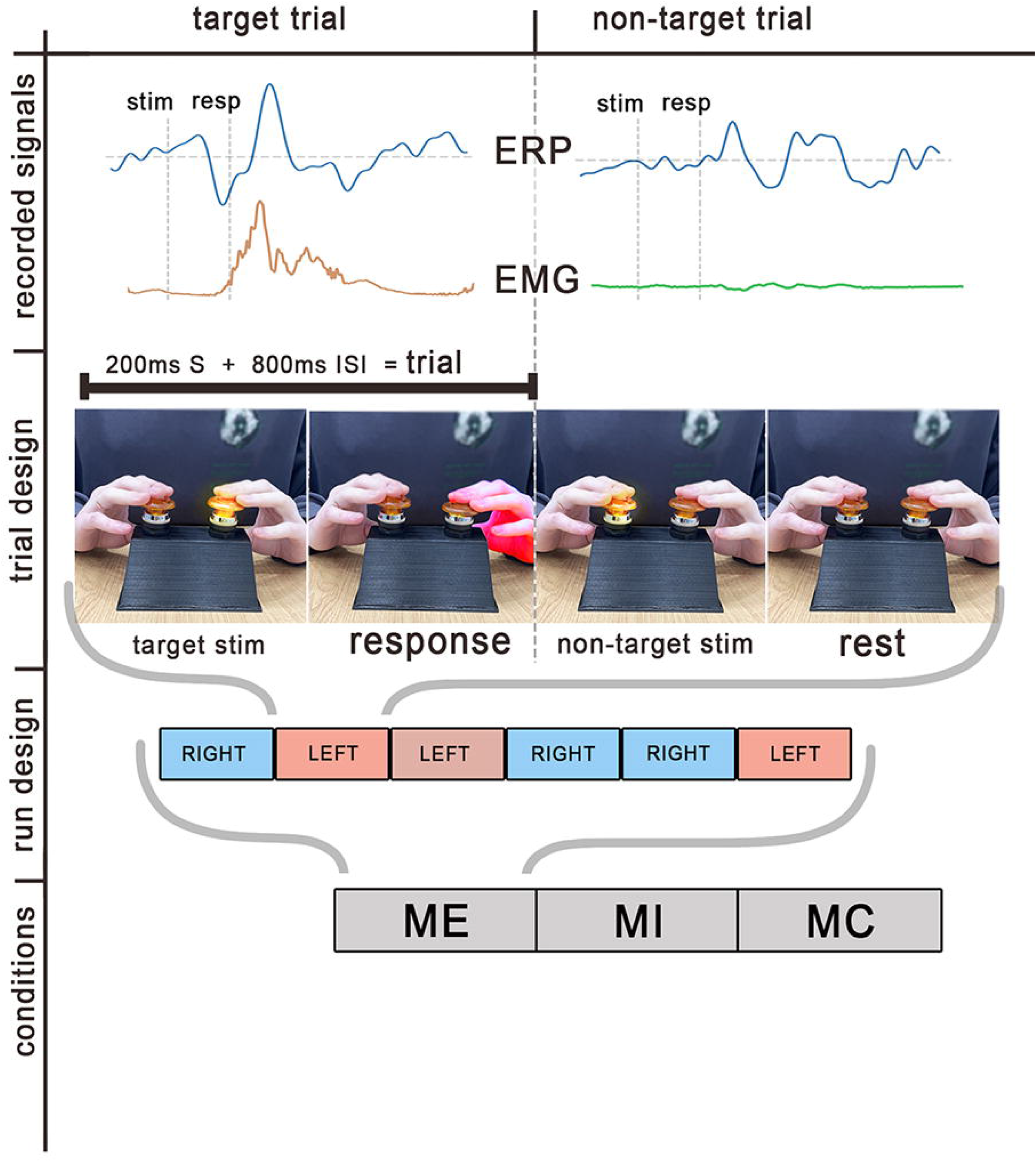
Experimental design. Within each experimental session, three tasks were run: motor execution (ME), motor imagery (MI) and mental counting (MC). Each run started with a command indicating which button was the target (RIGHT or LEFT). Participants focused on the target button highlights to which they responded by pressing the target button, imagery of button press or counting the number of highlights for the ME, MI and MC tasks, respectively. Each run contains 30 trials of button highlights, which were target and non-target highlights in equal proportion. The flash duration was 200 ms (S) and the interstimulus interval (ISI) was 800 ms. EEG and EMG signals were recorded. Stimulus presentation time and response time were synched with signal recording. Stimulus-locked and response-locked ERPs were analyzed for each condition.

### Signal acquisition

EEG was recorded at 1,000 Hz using an NVX52 DC amplifier (MKS, Russia) with 22 Ag/AgCl passive electrodes in the following positions: *Fp1, Fp2, Fz, FC3, FCz, FC4, C5, C3, Cz, C4, C6, CP3, CPz, CP4, P3, Pz, P4, PO3, POz, PO4, O1, O2 (with correspondence to the international “10-10” system)*. The average of channels A1 and A2 was used as the reference. The electrode–skin impedance was kept below 20 kΩ. The stimulus presentation time and the button press timestamps were synched with signal acquisition. Two channels of surface EMG for the left-hand and right-hand *m. flexor digitorum superficialis* (FDS) were sampled in order to discard non-ME-trials and non-target trials where muscles were activated.

### ERP analysis

The raw EEG signal was bandpass filtered using a Butterworth 4th order filter with a frequency range of 1-15 Hz. Blinking artifacts were removed using independent component analysis (fastICA method). The preprocessed signal was epoched [-0.5 – 1] s after the visual stimulus onset, the mean voltage of the [-0.5 – 0] s period was used for the baseline correction. We call these epochs “preprocessed epochs”. Then the stimulus-locked averaging was used to compute target and non-target ERPs in each condition. Furthermore, to compare stimulus- and response-locked ERPs during the ME condition, averaging was performed for the data aligned on the button press. To visually compare the stimulus and response-aligned ERPs, the 298-ms shift was performed, which corresponded to accounting for the median reaction time calculated for all subjects.

To compute lateralized potentials across all experimental conditions, stimulus-locked epochs were utilized due to the lack of overt responses during the MI and MC tasks. Considering the contralateral predominance of both pre- and post-movement MRPs, the calculation of lateralized evoked potentials (LEPs) was performed using the full-length epochs and the double subtraction method introduced by Coles (1989). This method is widely used to assess lateralized readiness potentials as established in previous studies (Carrillo-de-la-Pena et al., 2006). In this method, at first lateralized potentials were obtained separately for each trial by subtracting the epochs derived from the ipsi- and contralateral central electrodes (C3 and C4, see Eq.1, *a* and *b*) with consequent averaging (Eq. 1, *c)*:

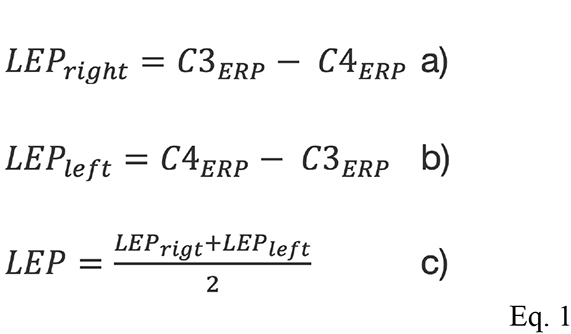

### Analysis of ME-related potentials

To analyze the stimulus-related action preparation and post-movement cortical activity in relation to the motor reaction time (RT), the single trial LEPs referred to the ME condition were obtained separately for each button press without the averaging (only Eq. 1, *a-b*). The same approach was used by Falkenstein et al. (2006). All participants’ LEPs waves were collected in one matrix and ranged by the ascending of the motor-response latency. The epochs with very slow responses (RT exceeds threshold 577 ms that is a computed Q3+1.5IQR value for all RT data) were rejected from the further analysis. The resulting matrix was visualized (Fig. 3). To reduce contamination of LEPs by oscillatory EEG components, a moving average with 10 ms window and one-timestamp steps was applied to each LEP. Amplitude, latency, and length of LRP were measured. LRP amplitude was measured as the peak negative LRP deflection within the time interval from visual stimulus onset till 80 ms after the response. LRP’s latency was defined as time of peak reaching. The length of the LRP was determined by measuring the width of the LRP deflection where the amplitude exceeds half of the peak amplitude (in absolute value). Additionally, we measured the latency and amplitude of RAP. The association between these parameters and the delay in reaction time was evaluated by calculating Pearson’s correlation coefficient across trials.

To directly estimate the differences in LEPs between the fastest and the slowest responses, the RT binning approach was applied (Roth et al., 1978; Poli et al., 2010). The approach consisted of ranking epochs based on their RTs and computing averages separately for different quartiles. The trials from the first quartile (Q1) corresponded to the fastest responses, the third quartile (Q3) represented slow responses. We didn’t use the epochs from the fourth quartile because as noted by Poli et al. (2010) these data correspond to noisy trials. Accordingly, two averages were calculated for all subjects: LEPs with the *fastest* responses and *slowest* responses.

To assess the significance of a lateralization of the LEPs components, ERPs from quartiles 1-3 were compared to the zero level that represented an absence of lateralization. We also computed LEPs for non-target responses and statistically estimated whether the lateralization appears in trials where no movements were present.

### Analysis of MI-related ERPs

For MI-related ERPs the late cognitive positive peak was analyzed, which is similar to RAP. Since alignment on response onset could not be done for MI trials, an alternative approach was implemented to estimate the ERPs, which consisted of latency shifting. Latency shifting is based on a trial-by-trial estimation of a particular ERP peak which is informative of the latency. Next, the estimated latencies are used to shift trials so that all peaks are collected at one time, i.e., resynchronized. This procedure corrects the latency jitter and improves the signal to noise ratio for various ERP components (Picton et al., 2000, Poli et al., 2010). However, accurate alignment may be affected by EEG fluctuations that introduce noise. To address this issue, we used a peak-picking approach proposed by Gratton et al. (1989) to realign ERPs. This procedure involved the application of a band-pass filter in the range of 0.5-2 Hz, followed by smoothing using a moving average procedure with a 20 ms window. The latency of the late positive peak of the MI-related positive peak was determined by identifying the maximum value within the 400-800 ms interval from stimulus onset. These determined latencies for each trial were then used to realign the unsmoothed epochs that were filtered with a wider band (“preprocessed epochs”) (Makeig and Onton, 2011; Ganin and Kaplan, 2022).

To evaluate inter-trial variability in the latency of the MI-related positive peak and RAP latency in ME task, the median absolute deviation (MAD) of peak latency was computed for each subject for the Cz lead (Ganin and Kaplan, 2022).

### Analysis of ERD

Following the preprocessing described above, a time-frequency analysis was performed. EEG signals were re-referenced to the common average reference (CAR) to enhance the spatial resolution of the signal. The Morlet wavelet transform was used for the analysis of time–frequency perturbations. We used a set of complex Morlet wavelets with variable number of cycles for different frequencies. The frequencies ranged from 5 to 25 Hz with 1-Hz steps, and the full-width at half-maximum (FWHM) was equal to 374 ms. The desynchronization value was calculated as the ratio of the signal power during the target epochs to the median of the signal power during the non-target epochs, which were restricted to the [0.5-1] s interval. The obtained values were converted to decibels (see Eq. 2). Negative and positive values corresponded to event-related desynchronization (ERD) and synchronization (ERS), respectively.

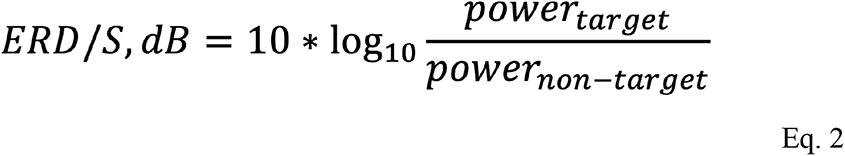

Medians were then calculated across all epochs to obtain ERD values for each of the experimental conditions (MI, ME, and MC). Our particular focus was on examining changes in β-power (15-25 Hz). We identified a subject-specific subrange within the β-band where ERD was predominantly expressed and calculated the median value within that particular subrange. The ERD values from channels C3 and C4 were then used for subsequent ANOVA and correlation analyses.

### Statistics

The Cluster-level statistical permutation test was used for the following comparisons: 1) whether the target and non-target LEPs significantly differed from zero (significant of the lateralization); 2) whether LEP amplitudes differed for fast versus slow responses. The permutation cluster-based test utilized in this study uses permutations and cluster-level correction to compute statistics that account for multiple comparisons. Specifically, a null distribution was generated by creating 100 000 random sets of permutations. Statistical significance was determined by evaluating the proportion of elements in the null distribution that exceeded the observed maximum cluster level. The F-threshold was automatically selected to correspond to a p-value of 0.01, which provided adequate control of the Type I error rate.

Kolmogorov–Smirnov tests were applied as tests for normality. A nonparametric Wilcoxon signed-rank test and parametric t-test were used for paired comparisons. In the ANOVA analysis, EEG desynchronization was assessed for three factors: condition (MI vs ME vs MC), hand (right vs left) and time (five consecutive 200-ms intervals starting with -100ms relative to stimulus onset). A Bonferroni correction was applied to correct for multiple repetition of the tests. Pearson’s correlation test was used to assess the relationship between variables. Specifically, we calculated the between-subject correlation between ERD scores and RAP peak.

## RESULTS

EEG responses to target stimuli differed from those to non-target stimuli in all conditions (Fig. 2, A). They consisted of the initial component P200 followed by negativity spanning the interval 200-400 ms after stimulus onset. Following these early responses, a late positive component developed in the ME and MI conditions. The characteristics of this component’s latency indicated that it was a potential time locked to the motor response, specifically RAP (and RAP-like potential during MI). The response-locked averaging of ME-related epochs confirmed the motor-response locking of this component (see Fig. 2, B). Indeed RAP amplitude was higher for the alignment on motor response compared to the alignment on the stimulus (Fig. 2, B). The RAP-peak was stronger in the hemisphere contralateral to the acting limb. The negative deflection observed in ME-trials (named as N2 in Fig. 2) was more prominent in the contralateral hemisphere, as well. As its peak was more negative in stimulus-locked averaging, topography maps were computed for this averaging approach. Since ME-related N2 is more prominent contralaterally, it is consistent with the definition of lateralized negative readiness potential.

**Figure 2.**
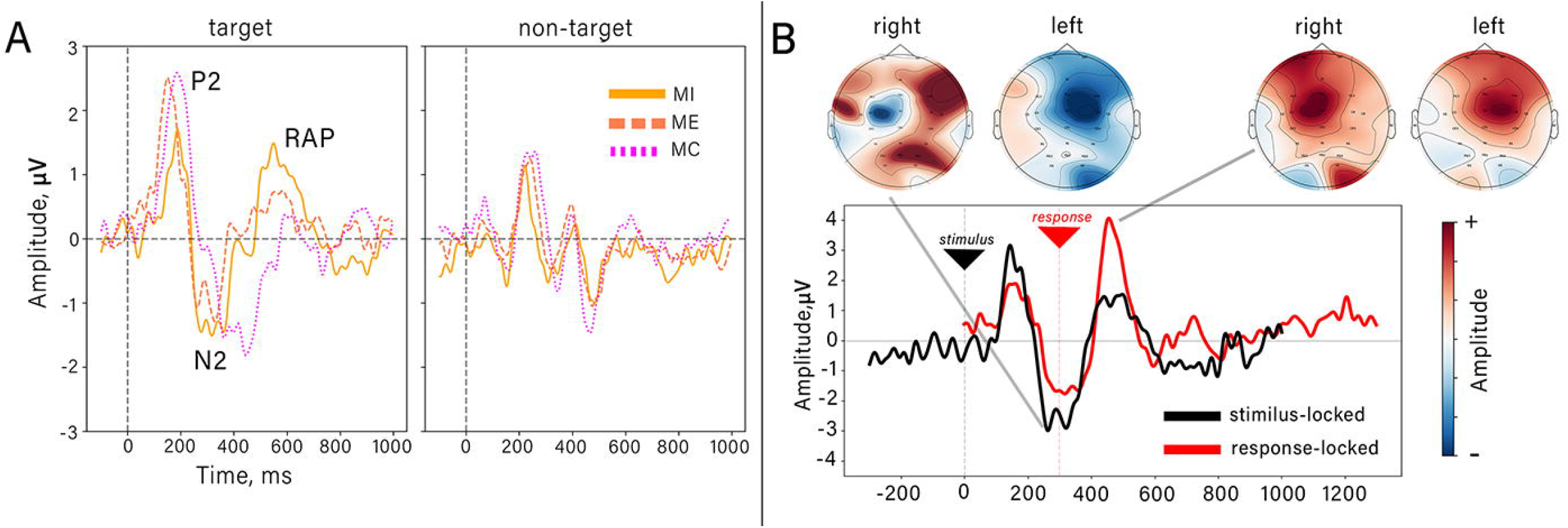
**A.** Stimulus-locked ERPs in the target (right) and non-target (left) trials. **B.** Stimulus-locked (black) and response-locked (red) averaged ME-related ERPs plotted in the same graph. Additionally, topographical distribution is shown of peak amplitude of the corresponding motor-related potentials separately for left- and right-responses. Red shapes indicate positive values, and blue indicate negative.

### ME related LEPs and the reaction time

Figure 3 shows the across-participants LEPs from ME-trials for different reaction times. In this plot, color-coded LEPs are ordered according to the reaction time. Time zero is visual-stimulus onset, and motor response onset is represented by a bold dashed line. Two LEP components were apparently observed. The first component was a negative LRP that started ∼200 ms prior to the motor response and disappeared after or simultaneously with the motor response. Positive correlations were found between the reaction time and latency of the LRP peaks (r=0.60,p=5.2e-110) and between the reaction time and LRP duration (r=0.63,p=1.47e-125). No correlation was found between LRP amplitude and RT (see Fig. 3, B). While the LRP onset was rather time-locked to the stimulus, its duration extended to the motor response, i.e. the LRP lasted until the movement. This effect is clearly seen in Figure 3, C, where the ERPs are shown for the fastest and slowest quartiles of RTs. It can be seen that for the slowest RTs, the negative deflection before movement onset was longer than for the fastest RTs. Furthermore, the amplitude of the LRP is significantly lower for the slowest RTs (p < 0.01 for a cluster spanning the 150-210 ms range), even though no significant correlation was found.

**Figure 3.**
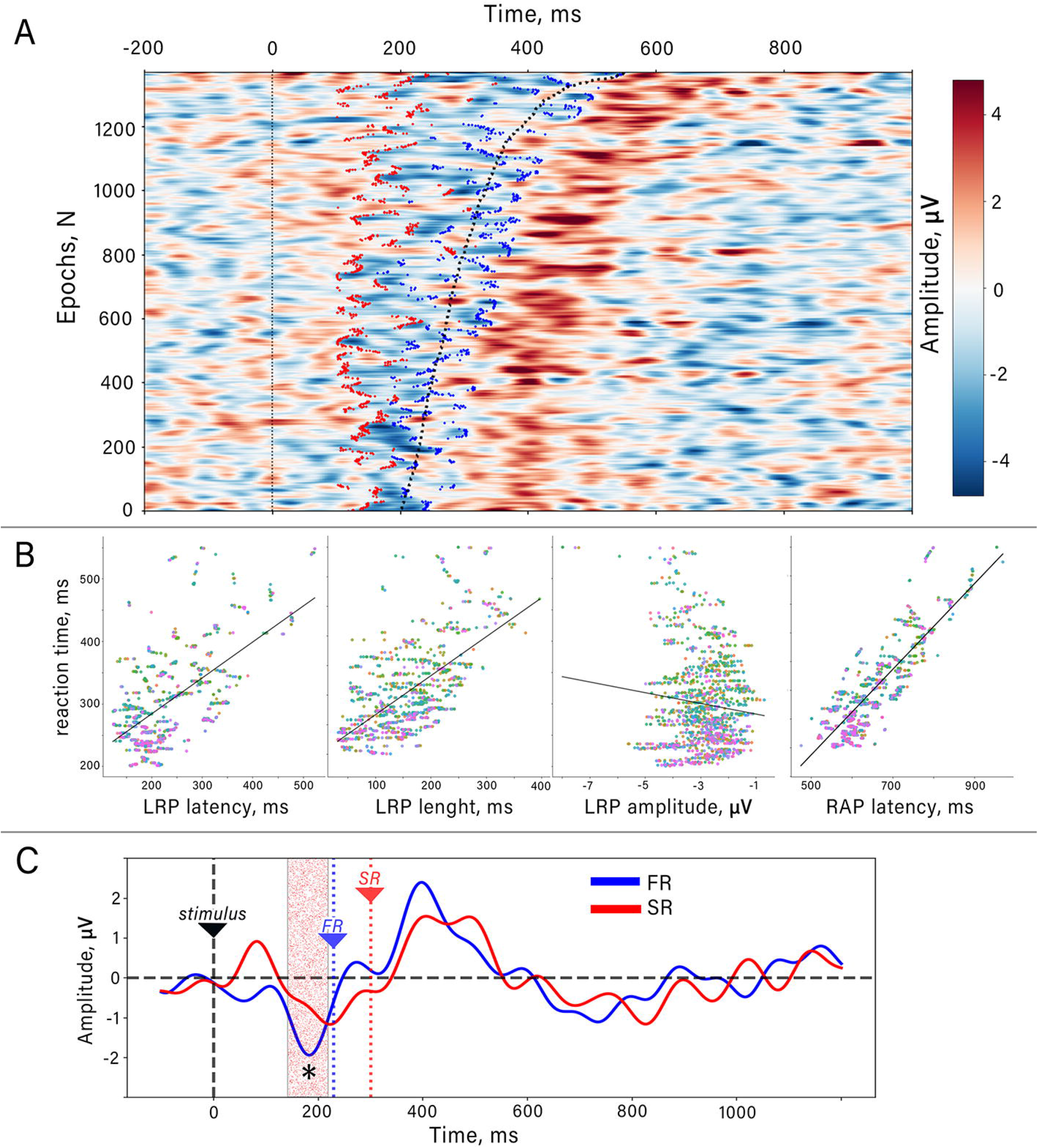
**A.** LEPs for all ME epochs in all participants sorted by reaction time (bold dashed line). Red and blue dots limit the LRP’s onset and offset determined as the period where LRP deflection exceeded 50% of maximal amplitude. **B.** Correlations between response reaction time and LEP parameters (from left to right: LRP’s latency, LRP’s length, LRP’s amplitude, RAP’s latency. Dots color indicates different participants. **C.** LEPs for the fastest (FR) and slow (SR) responses averaged across subjects. Bold black dashed line indicates stimulus onset, blue and red dashed lines indicate median response time for the fast and slow LRP’s respectively. Filled area marked by an asterisk indicates the temporal cluster where significant differences were revealed (p < 0.01, nonparametric cluster-based permutation test).

The positive deflection of the EEG potential that followed the LRP (see Fig. 3, A) matched the definition of RAP (Vaughan, 1969). The RAP onset was clearly time-locked to the motor response onset which it lagged by ∼200 ms. A significant correlation was found between the RAP latency with respect to stimulus onset and motor reaction time (r=0.83, p<0.0001, see Fig. 3, B).

Figure 4 shows the subject averages of the stimulus-locked LEPs for the target (blue line) and non-target (gray line) trials where no muscle activity was observed. For both target- and non-target trials, cluster-based permutation testing revealed significant (p<0.001 and p<0.01 for targets and non-targets, respectively) difference from the zero level. The temporal clusters revealed with this method matched the LRP latency range corresponding to the premovement period demarcated by the EMG onset. A slight negative deflection was found for non-target trials, which had a lower amplitude compared to the target trial LRP. This deflection returned to baseline by the time the EMG response began to develop in the target trials.

**Figure 4.**
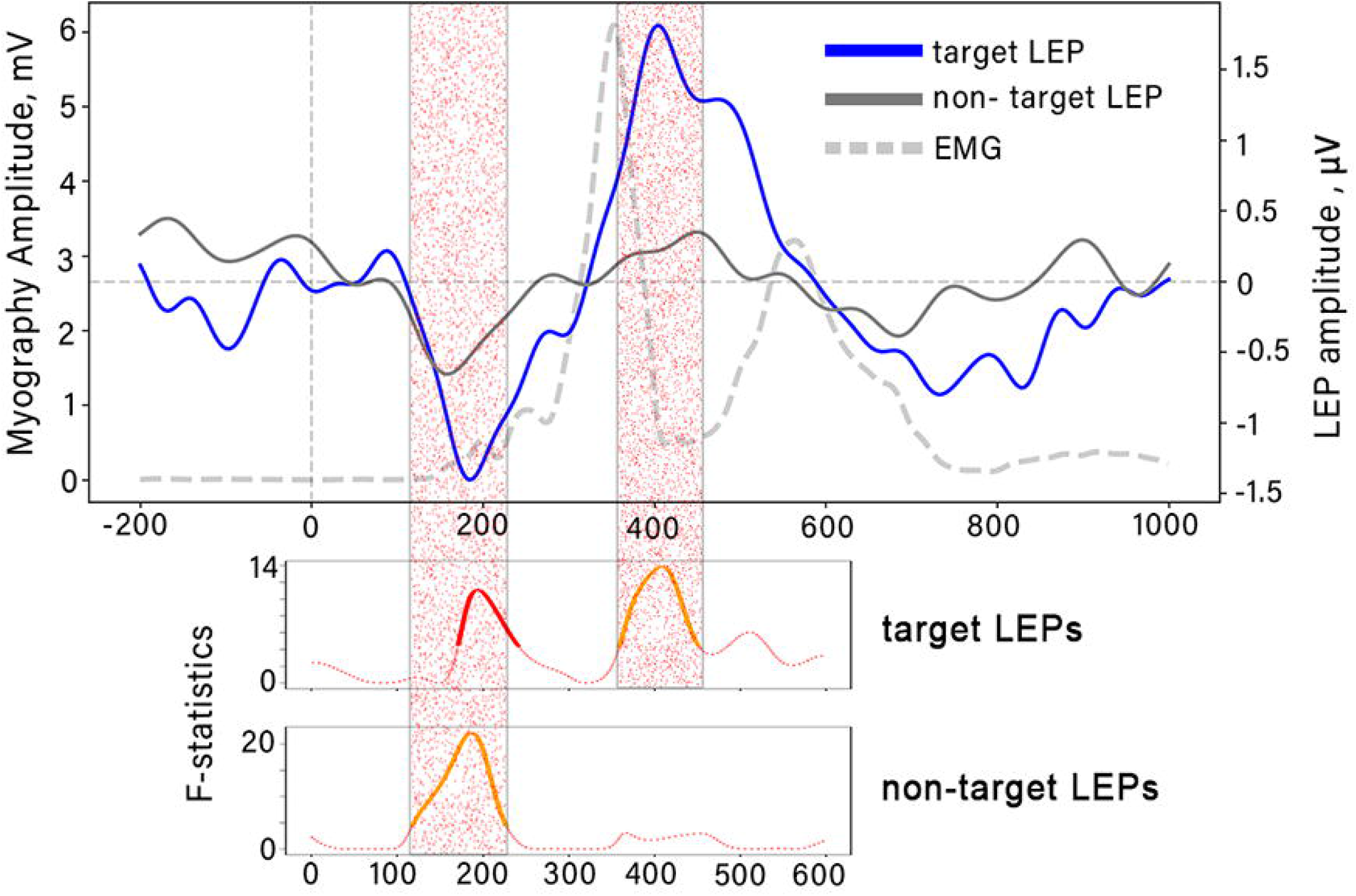
**A.** Stimulus locked across-subject average LEPs for target (blue line) and non-target (gray line) ME-trials. Dashed transparent gray line is an averaged EMG response corresponding to target trials. **B.** Results of the nonparametric cluster-based permutation tests indicate the significant differences between LEPs and zero. F-statistics is shown. The red color indicates temporal clusters with p<0.001, orange line indicates p<0.01, the dotted line corresponds to insignificant differences.

Target-trial LEPs were characterized by a significant contralateral positive RAP (p<0.01) that reached maximum at the time of maximal muscle activity and returned to the baseline when the button was released.

### Motor imagery-related ERPs

In the MI condition, target ERPs had a positive component spanning the 500-700 ms interval relative to stimulus onset. This interval overlapped the latency interval of the RAP component observed in the ME trials. We refer to this component as the RAP-like potential because it was maximal over the sensorimotor areas (i.e., channel Cz). Unlike movement-evoked RAP, the RAP-like potential has no hemispheric predominance, so the lateralized potential in MI trials was not significantly different from zero (see Fig. 5). Notably, the N2 component was not lateralized during MI, regardless of the hand whose movements were imagined. No lateralized component was found for mental counting trials, as well.

**Figure 5.**
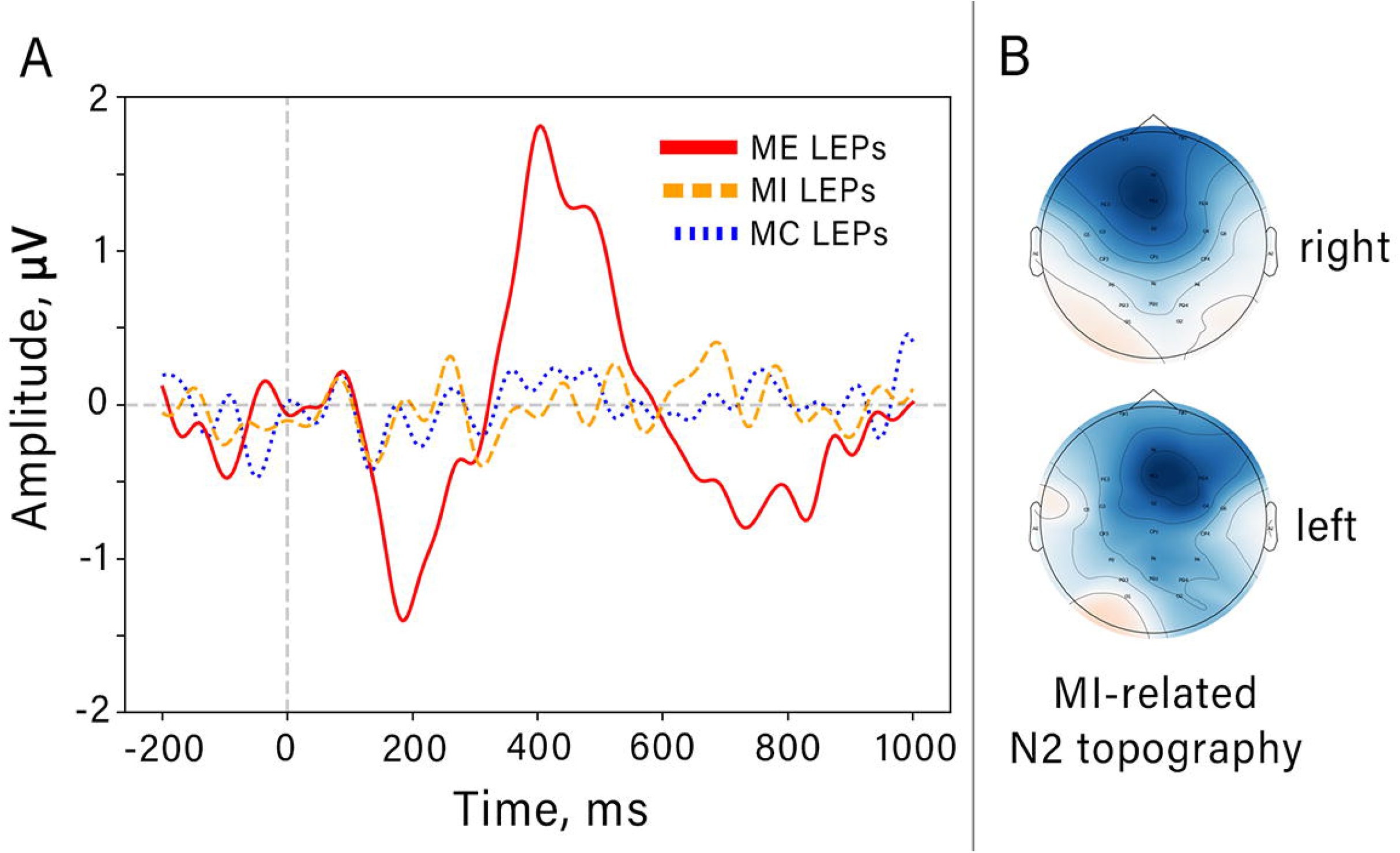
**A.** Lateralized evoked potential (LEPs) in three experimental conditions: MC, MI and ME. **B.**Topographic distribution of the N2 components in MI-related potentials for “left” and “right” trials.

Because the late positive peak observed during MI is the mental response-locked, its amplitude found with stimulus-locked averaging must be incorrectly defined due to latency jitter. To test this hypothesis, we performed ERP-locked latency jitter adjustment where we estimated the variability of RAP-like latency. Figure 6 shows the data for MI target epochs for all participants, ordered by estimated RAP-like latency. This analysis showed that the latency of the early positive component P2 was locked to the visual stimulus onset. The negative wave N2 started ∼250 ms after the stimulus and lasted until the RAP-like positivity onset whose latency ranged from 400 to 800 ms. Figure 6, B shows an across-subject average ERPs for the MI condition. The peak-locked lining procedure did not result in a significant change of the amplitude of the RAP-like component. A comparison of the median deviation of RAP latency in ME condition with the RAP-like component. revealed a significantly higher variability of the movement-related RAP (Wilcoxon test, W=1, p=0.0019).

**Figure 6.**
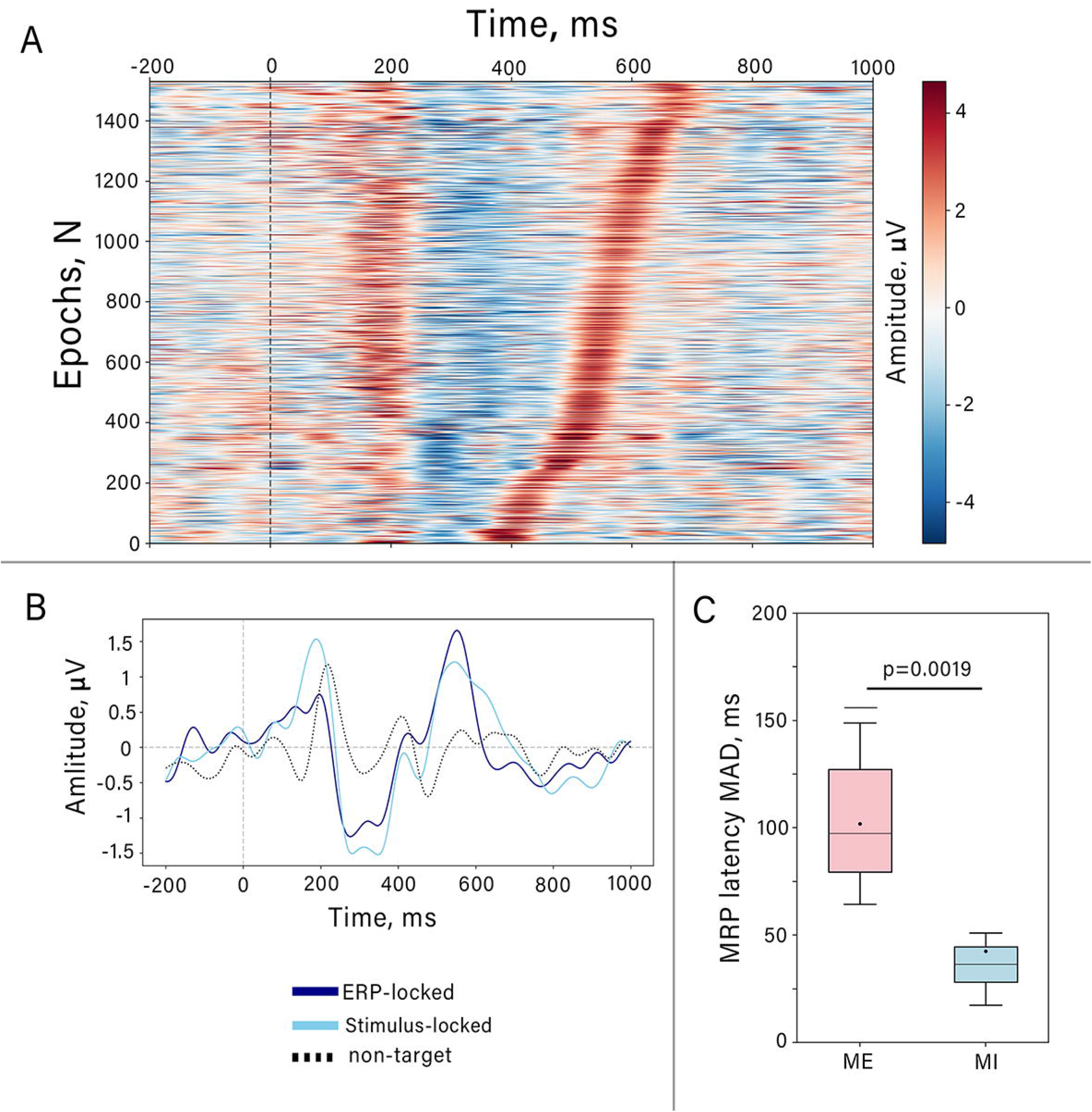
**A.** The epochs related to target MI-trials in all participants sorted by ascending of determined RAP-like latency time. Dashed line indicated stimulus onset. **B.** Grand averaged MI-related ERPs in Cz channel before (dark line) and after (light line) latency adjustment. **C.** Comparison between median absolute deviation of RAP and RAP-like components in motor execution (ME) and motor imagery (MI) trials.

### β-ERD analysis

The analysis of event related desynchronisation in the β-band was carried out to assess the dynamics of sensorimotor cortical areas activity in ME, MI and MC conditions. Short-lasting contralateral ERDs were observed during both MI and ME. Figure 7, A shows the grand-averaged time-frequency dynamics for the right-MI and left-MI conditions; channels C3 and C4 are illustrated. The most prominent ERD occurred in the β-frequency range (15-25 Hz) and it peaked 500-600 ms after visual stimulus onset. Conversely, mental counting did not reduce the amplitude of β activity (see Fig. 7, B). Three-way ANOVA revealed significant main effects for the condition (MI vs ME vs MC, F=30.81, df=2, np2=0.11; p<0.0001) and time (F=4.05, df=4, np2=0.05, p=0.003) factors but not for the hand factor. In addition, significant interactions between condition and time were found (F=6.62; df=8, np2=0.097 p<0.001). Post-hoc comparisons conducted for the data from 4th time-interval (500-700 ms range) confirmed a significantly greater ERD amplitude in MI and ME conditions compared to MC (W=54, p=0.000008 for ME and W=31, p=0.00001 for MI; see Fig. 7, B). However, no difference in the β-ERD values were found for the comparison between the ME and MI trials (W=162, p=0.056).

**Figure 7.**
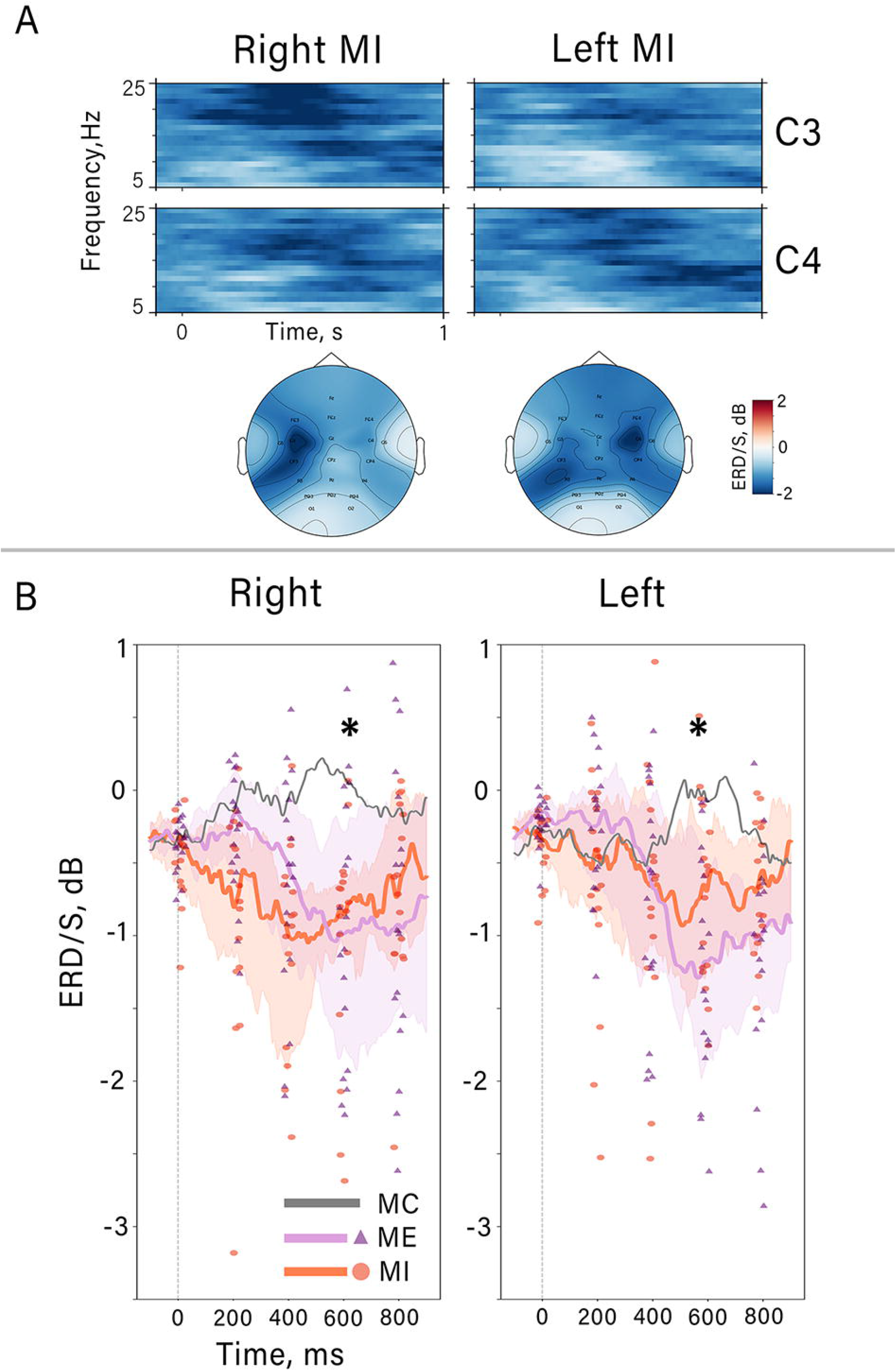
**A.** Median across subjects showing time-frequency dynamics of MI target trials in C3 and C4 channels and corresponding topography distribution (bottom). **B.** Average time courses of β-ERD for all experimental conditions in C3 channel for right-responses and C4 for left-responses. Lines are median, colored shapes indicate interquartile range. Dots are individual ERD values for all subjects averaged in 200 ms time-interval. Bold asterisks indicate the time-interval where post-hoc tests revealed significant differences.

Figure 8 shows the presence of negative correlation between the amplitude of β oscillations and the amplitude of the RAP-like peak. Pearson’s test revealed moderate significant correlation (r=-0.67, p=0.0035) for the comparison of RAP-like amplitude measured for channel Cz and peak MI-related β-ERD amplitude for the contralateral channel (since the hand factor did not affect the ERD dynamics, here we combined the data for the right and left hand).

**Figure 8.**
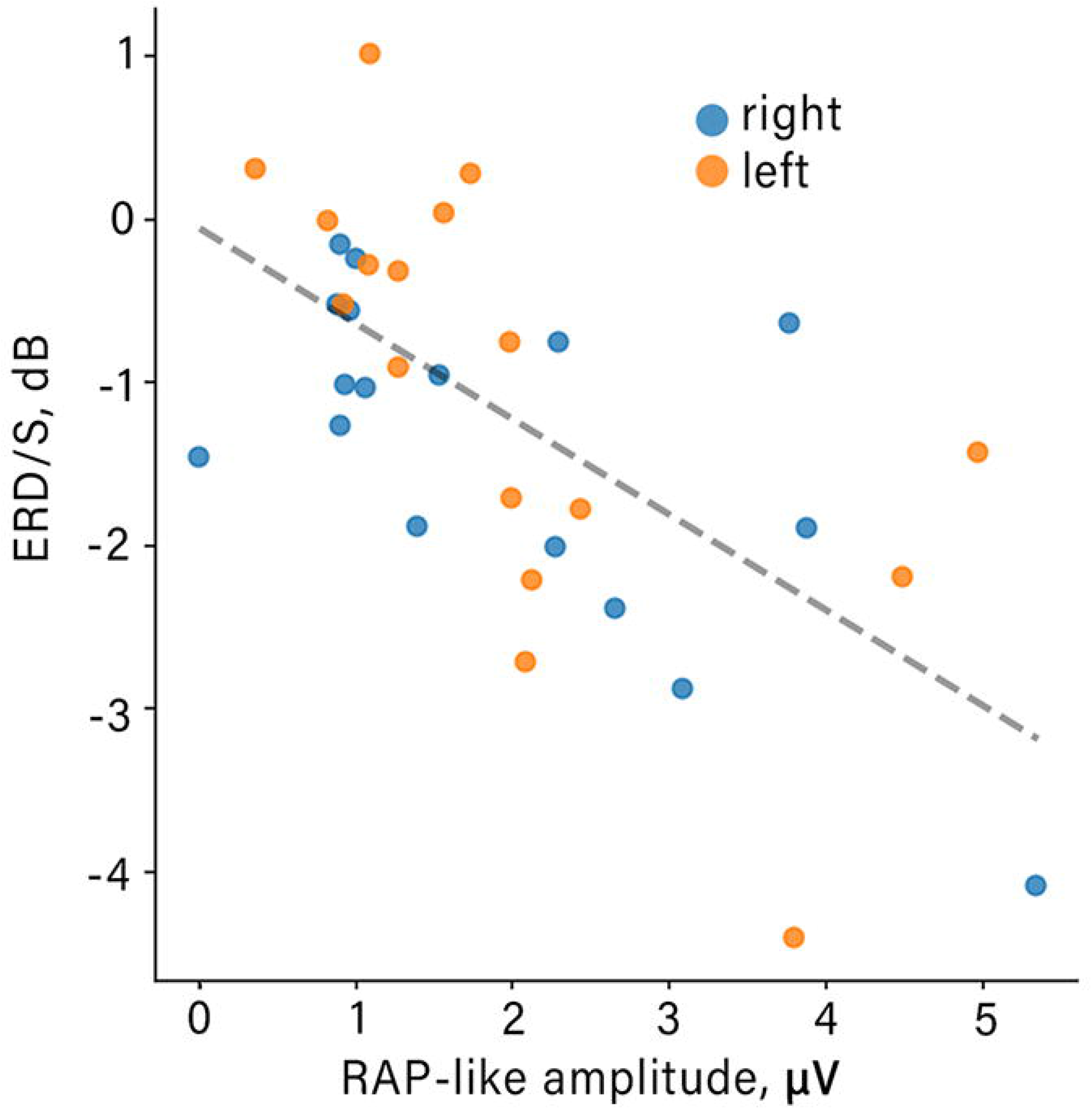
Correlations between β-ERD value and RAP-like amplitude for left and right MI-responses.

## DISCUSSION

In this study, we examined EEG responses to target and nontarget visual stimuli for three instructions: mentally counting the target stimuli (MC), responding to them with button press (ME), and imagining button-press responses (MI). Lateralized ERPs, as markers of limb-specific sensorimotor area activation, were assessed across these conditions. Additionally, an analysis was performed of the relationship between the timing of different MRP components and reaction time. These analyses clarified the process of visuomotor transformation, where a visual stimulus triggers different stages of neural processing in the sensorimotor cortex related to action preparation and execution.

### The relationship between LEP components and reaction time in terms of motor accumulation theory of action

Premovement potential with a pronounced lateralization was clear only in ME trials. Its amplitude was maximal over frontocentral areas in the hemisphere contralateral to the active limb. This kind of EEG potential previously was called LRP. LRP develops in the motor cortex during movements preparation, typically about 200ms prior the EMG onset (Böcker et al., 1994). At the time of LRP development, subjects are usually aware of their decision to act (Trevena and Miller, 2002; Neafsey et al., 2021), so the LRP is thought to reflect primary motor cortex (M1) activation during the final stage of action preparation. LRP was described as a ballistic stage where the motor gate has opened and the signal from the brain rushes to spinal motoneurons (Logan et al., 1984; Schultze-Kraft et al., 2016) and it becomes impossible for the nervous system to veto the upcoming movement after LRP onset (Schultze-Kraft et al., 2016).

While the LRP is commonly characterized as a component locked by movement onset, it also provides insight into the neuronal processing of the stimulus and its subsequent translation into movement. This is evident from the relatively narrow peak of the stimulus-locked LRP, suggesting the involvement of stimulus-related processing stages in shaping the LRP response (Hohlefeld et al., 2011). Accordingly, a better description of LRP is in terms of stimulus-response mapping process rather than merely motor preparation. In support of this view, LRP shape and latency depend on the stage of perceptual categorizing of the stimulus (*e.g*., linking the stimulus to the limb that has to move) followed by the stage of motor preparation and execution (Hackley and Valle-Inclan, 1998). Luck et al. (2009) provided further evidence linking the LRP to stimulus processing. They observed slower reaction times in individuals with schizophrenia compared to healthy participants. Although a slight delay in the latency of the response-locked LRP was observed in the patient group, it does not fully explain the observed RT slowing. On the other hand, a study by Saville et al. (2015), which examined the correlation between reaction time (RT) and LRP latency in patients with attention deficit hyperactivity disorder (ADHD) characterized by highly variable RTs, proposed that altered motor cortex excitability and impaired resolution of interhemispheric competition, as well as increased neural noise, contribute to this relationship. The drift diffusion model clarifies these relationships further. According to this model, a stimulus being analyzed during decision making triggers a noisy evidence-accumulation process where the activity of neural circuitry drifts toward a decision threshold (Van Vugt et al., 2014; Hanks et al., 2015). Based on such a model, Van Vugt et al. (2014) proposed that the early part of the LRP reflects the accumulation of evidence, while the later phase is a self-maintaining ballistic formed by a motor network that has passed a decision threshold (Van Vugt et al., 2014).

In the present study, we found further evidence that the LRP represents a visuomotor transformation by examining single-trial lateralized EEG responses in relation to reaction time. We found a moderate correlation between LRP timing and RT in the paradigm where evidence accumulation consisted of dissociating targets from non-targets, followed by choosing the acting hand. However, the results obtained indicate that the early phase of the LRP, which reflects the accumulation process, is not sufficient to fully explain the observed delays in RT. This suggests that the response delays are also influenced by the excitability of a "threshold layer" that varies across trials, contrary to the previous hypothesis proposed by Van Vugt et al. (2014). This variation apparently contributed to the LRP shape as not only LRP shifted with the RT but its duration increased with the increases in RT. Similar results were obtained by Sangals et al. (2002) who reported shortening in LRPs for faster motor responses. Furthermore, our observation that LRP duration was correlated with RTs, but not its peak amplitude, is consistent with the model in which the slope of increasing neural activity (rather than the peak amplitude) is important for reaching a fixed threshold and triggering a response (Hanes and Schall, 1996; Sangals et al., 2002). However, in a more sensitive analysis, we found a relationship between RT and LRP amplitude. We split the trials into two groups, one with the slowest RTs and the other with the fastest RTs, and found greater LRP amplitude in the latter. Thus, a higher rate of LRP rise was also associated with a higher value of LRP peak. Regarding the reason why some responses were slower than others, we could speculate that fluctuations in motor cortex excitability (Misirlisoy and Haggard, 2014; Aksiotis et al., 2022) may have played a role. This suggestion is supported by the observations of Yilmaz et al. (2015) and Roushdy et al. (2022) who demonstrated alterations in premovement cortical potentials in post-stroke hemiparetic patients at both acute and chronic stages, which they related to interhemispheric inhibition of the affected sensorimotor cortex by the intact contralateral hemisphere. Additionally, Churchland et al. (2010) proposed that premovement activity sets up initial conditions for the subsequent cortical dynamics that subserve movement execution (Churchland et al., 2010).

Variations in LRP duration are of particular interest for understanding the underlying visuomotor transformation. In the future, these variations could be further investigated by enriching our paradigm with more complex stimulus-response compatibility (Brass et al., 2001) and eliciting involuntary movements by coupling motor imagery with action observation (Colton et al., 2018). In a real-time scenario, EEG activity could be monitored for features that affect RT, such as β power (Paluch et al., 2021; Aksiotis et al., 2022). Additionally, motor cortex readiness to execute a movement is known to be influenced by internal factors (Kunzendorf et al., 2019; Al et al., 2021) and stochastic processes (Murakami et al., 2014; Murakami et al., 2017; Aflalo et al., 2022). In particular, an increase in LRP duration associated with RT slowing has been documented in the elderly population (Falkenstein et al., 2006), which on the one hand suggests a deterioration of different stages of sensorimotor transformation with aging, and on the other hand makes BCI paradigms such as the one proposed in (Syrov et al., 2022) a useful tool for monitoring and possibly treating such age-related changes.

### Non-target LRPs

Curiously, even though participants in these experiments focused their attention on the target stimuli and ignored the distracting non-targets, LRPs were observed also in response to non-targets. In the ME condition, these non-target LRPs were small and of shorter duration (Fig. 4). Although target and non-target LRPs started at the same time relative to the stimulus, non-target LRPs returned to baseline at the time that matched EMG onset in the target trials. This was also the time when LPRs reached their peak value in the target trials. Notably, the sharp rise in EMG continued after the target LRP faded. The presence of non-target LRPs agrees with the model of decision-making where sensory information is evaluated in the context of the instructed motor task and a veto decision results from this evaluation. Previously, Carrillo-de-la-Peña et al. (2006) observed double-peak LRPs. They proposed the first peak could have represented the selection of the hand prior to the final decision on whether or not to initiate a movement.

Taken together, our results on the target and non-target LRPs provide clues on how cortical visuomotor circuitry propagates visual information and converts it into motor execution commands when appropriate. We think that it would be reasonable to subdivide the LRP into an initial phase with the timing corresponding to the duration of non-target LRPs corresponding to the evaluation of visual information and the late phase corresponding to motor preparation (or cancellation for non-target trials). This is not to say that cortical motor areas stay silent when the stimulus is evaluated.

On the contrary, they are activated during this process simply because they are part of the stimulus-response mapping circuitry and need to prepare their state for the possibility of subsequent movement initiation (Van Vugt et al., 2014). Thus, the mere presence of an LRP does not necessarily indicate an imminent motor response, nor does it directly signify the occurrence of action awareness. Rather, the presence of LRPs in non-target trials could be interpreted as an early preparation of a movement that was later interrupted. In a paradigm similar to our non-target trials, Galdo-Alvarez et al., (2016) showed LRPs were lower in no-go trials, where participants successfully stopped movement preparation. Conversely, LRPs were stronger in the no-go trials where the participants failed to stop movement. These observations show that LRPs result in movements only if they are strong enough. In the present study, in only 0.12% of non-target trials EMGs were detected, so we do not have a sufficient amount of data to directly compare our results with those of Galdo-Alvarez et al., (2016). Yet, our results could be explained in a similar way. The motor circuitry could have been in a state of persistence readiness to react as fast as possible, so any visual stimulus started a visuomotor transformation-like process where motor preparation and execution was allowed to fully develop only in the target trials. This type of neural processing is similar to the previously described central command-relieved vetoing (Schultze-Kraft et al., 2016), and the ability to successfully interrupt an action in preparation requires motor cortical activity to be low enough (Misirlisoy et al., 2014).

### An absence of lateralized potential in the MI and MC tasks

Although lateralization of cortical potentials was very clear in the ME task, we did not observe lateralization for MI and MC even though we observed N2 potentials that had the same latency as the LRPs in the ME condition. Thus, the N2 wave was not lateralized during MI even though the subjects imagined movements performed by the hand adjacent to the target button. This result disagrees with several previous reports where LRPs were found during MI that were weaker (Galdo-Alvarez et al., 2016) or even comparable (Caldara et al., 2004) to LRP amplitude in real execution. Furthermore, an inversion of LRP polarity was observed when foot movements were imagined (Carrillo-de-la-Pena et al., 2006). Although these previous results appear to support high similarity of cortical potentials associated with the executed and imagined movements, our present findings show that this conclusion cannot be extended to all experimental conditions. Similar to our present results, Hohlefeld et al. (2011) did not find lateralization of cortical potentials for imagened and quasi-movements of the hand (Hohlefeld et al., 2011). They suggested that sensorimotor activation was weak under these conditions with covert movements. Such weak activity without lateralization could take place before response preparation becomes effector (hemispherically) specific (Saville et al., 2015). A somewhat similar turn of a non-lateralized potential into a lateralized activity was demonstrated by Carrillo-de-la-Peña et al. (2008), who observed this effect in both ME and MI conditions in a paradigm where an imperative stimulus evoked a non-lateralized negativity that became lateralized after the presentation of a cue instructing which hand should perform an action or be imagined doing so.

It should be noted that we used a block design where subjects did not change the working hand in a block of trials, so there was no trial by trial requirement to select the type of motor response anew. These settings could explain the absence of lateralized cortical potentials in the MI condition because, as suggested by Carrillo-de-la-Peña et al. (2008), MI exhibits activation in the M1 analogous to actual execution specifically during the working hand selection phase, but not during other processing stages.

Despite the absence of lateralized responses, significant β-ERD occurred during MI trials but not during MC trials. β-ERD developed during ME trials, as well. Assuming that β-ERD is a sign of increased motor cortical activity (Yousry et al., 1997) it is reasonable to suggest that such activation reflected the presence of a motor component in the ME and MI conditions. In particular, our observations on the differences in ERD in the MI and ME conditions are consistent with the previous report by Hohlefeld et al. (2011), suggesting that although there are similarities between these two conditions as predicted by simulation theory (Jeannerod, 2001), imagining actions and executing them are enabled by separate states of cortical motor circuits. Specifically, MI and ME conditions differ in the strategy on how to react to external stimuli where sensorimotor oscillations could have a role. According to Jeannerod’s theory (Di Rienzo et al., 2014), MI does not lead to actual motor performance due to inhibition of motor commands. However, our observation of non-target LRPs in ME trials, but not in MI trials, supports the idea that prior strategy, whether for a mental or physical response, influences the activation of M1. This suggests that preparation for an imagery button press does not require readiness of the motor system, since no actual motor response is required. We hypothesize that prior strategy control may occur via a highly hierarchical structure, similar to the ‘response layer’ proposed in the drift-diffusion model (Van Vugt et al., 2014). In contrast to the prevailing view that inhibition of execution in MI responses occurs via motor network suppression, we propose that the underlying accumulator layers (e.g., parietal areas) serve as targets for inhibition. This mechanism effectively prevents stimulus-response mapping processes and explains the absence of lateralized readiness potentials (LRPs) in MI compared to non-target motor execution (ME) trials. The presence of a downregulating command ("how to react") resulted in non-target LRPs that were incongruent with the stimulus but had a polarity consistent with the target hand. For example, when the right button was the target, non-target LRPs showed the same lateralization as in target trials, regardless of whether they were elicited by left button flashes. Consequently, the polarity of non-target LRPs does not show stimulus-response compatibility (Berlucchi et al., 1977; Holländer et al., 2011), but is consistent with the "which hand to respond’’ strategy determined by the prior instruction (also referred to as the "precuing effect" in Sangals et al., (2002).

Finally, previous experiments on single-unit responses in monkeys are useful for the interpretation of our present results. In a classical study, Georgopoulos et al. (1982) observed neuronal discharges in M1 that occurred ∼80 ms prior to EMG onset in the arm muscles, which corresponded to 100–150 ms after the visual cue. Ifft et al. (2012) and Lebedev et al. (1994) reported similar patterns of premovement activity in M1 and S1 elicited by the presentation of visual stimuli pointing in the direction of required action. The timing of these neuronal patterns was similar to the timing of LRPs that we observed here. Therefore we could speculate that our results could be reproduced in single unit-recording experiments in the future, perhaps in human participants because only humans can easily perform tasks like MI and MC.

### RAP and RAP-like potentials

Numerous studies have already explored the temporal relationship between the late ERP components and movements (Verleger et al., 2006; Saville et al., 2012; Verleger et al., 2014; Berchicci et al., 2016). Consistent with previous literature, we observed the development of a contralateral positive motor-related potential after movement onset. This potential was mostly distinct and strong when the EEG traces were aligned to the movement onset before averaging, and the latency with respect to the movement onset was ∼200 ms. This cortical potential is well known as re-afferent potential or movement-monitoring potential, and it has been suggested to originate in the precentral regions (Caldara et al., 2004) and somatosensory cortex (Bötzel et al., 1997). RAP is related to controlling movement execution while processing somatosensory feedback (Bötzel et al., 1997, Berchicci et al., 2016); therefore the synchrony between movement onset and RAP timing is rather obvious.

In our experiments, imagery of pressing a button led to the appearance of positive central-localized potential that had the same latency as the RAP in ME condition. However, the RAP-like MI-related peak was not lateralized. It was suggested that the amplitude of MRP’s components is proportional to the number of motor units recruited for the performance of a movement (De Morree et al., 2012; Sosnowska et al., 2021). As muscle activity is absent during MI, the presence of RAP-like in this condition was somewhat surprising. Several explanations for this effect could be proposed. First, MI-related positivity could have been a late P3 evoked by the target stimulus, however this potential was not presented in MC condition despite the necessity to process the target information during MC (Mertens et al., 1997). Secondly, our surface EMG could have been not sufficiently sensitive to detect EMGs in deeply located muscles and slight (subliminal) muscle twitches (Sosnowska et al., 2021). However, a study using ultrasound imaging to record deep muscles observed a RAP-like component during motor imagery without any discernible muscle twitching (Sosnowska et al., 2021). The third potential explanation could be in terms of emulation theory (Grush, 2004) which elaborates on the models of the body and environment constructed by the brain to control movements and adapt them to the environment. Sensorimotor efference copies of movements provide expectations of the sensory feedback which are then compared with the afferent inputs that occur during the movement.Such emulation could be run explicitly during MI (Sadato and Naito, 2004; Ridderinkhof et al., 2015), and then RAP-like potential could reflect the activation of cortical areas involved in this emulation. The absence of cortical areas responsible for executing actual motor acts in the cortical substrate of motor emulation results in the non-lateralization of the RAP-like component, similar to the behavior of the sensorimotor ERD, which shows greater lateralization during physical movement compared to motor imagery (Nikulin et al., 2008). Accordingly, the MI-related positive component can be viewed as a surrogate part of the normal process of predicting movement-related reafference that lacks the real sensory feedback processing. Given the reliance on afferent input delay, the comparison phase should show increased variability in the latency of RAP during ME compared to the entirely mentally generated RAP-like peak in MI trials. The greater median absolute deviation in RAP latency jitter compared to RAP-like peak provides empirical support for these theoretical speculations.

### β-ERD in MI and ME conditions

We found significant β ERD in ME and MI trials, but mental counting did not result in ERD exhibit. For the MI condition, we observed a gradual decrease in β amplitude for 500-600 ms from target onset. Such β-ERD is consistent with previous MI studies (Pfurtscheller et al., 2008), where it has been considered as a marker of activity in precentral cortical areas (Neuper et al., 2006; Babiloni et al., 2016). Revealed here β-ERD in MI trials can be also interpreted in terms of the emulation theory. Considering the potential connection between the RAP-like component and action emulation processes, as well as the β-ERD induced by motor imagery reflecting activation of the sensorimotor cortical substrate for movement emulation, it is plausible to explore potential correlations between these markers. Despite contemporary hypotheses proposing a common source for ME-related ERD and RAP, we did not uncover compelling evidence of a clear correlation between these markers. Our analysis of the across-subject correlation between β-ERD amplitude and late positive ERP component revealed significance correlation only for MI but not for ME. This result could be explained by the involvement of additional, reafference-related components during ME, which complicate the relationship between ERD and RAP.

The revealed correlation contributes to the relationships between MI-related positive peaks and sensorimotor activation, providing the use of MI-related ERPs to be useful in the design of BCIs. In summary, our goal was to carefully trace the temporal dynamics of the neural mechanisms involved in motor preparation and motor imagery. We achieved this by conducting a comprehensive analysis of EEG responses within an experimental paradigm of visuomotor transformation. By intricately examining the dynamic variations of various components of the MRPs in relation to reaction time, our study has detailed hidden stages of action preparation. These findings have remarkable implications for the practical use of the studied MRPs components as invaluable tools in the study and diagnosis of motor impairments.

### Limitations

A limitation of our study is that we did not perform comprehensive assessments of participants’ handedness. This lack of detailed testing prevents us from assessing hemispheric asymmetry, which is known to be relevant to motor control (Schmitz et al., 2019). It is worth noting, however, that a study by Schmitz et al. (2019) reported that handedness primarily affected LRP amplitude rather than its latency. Nevertheless, we intend to include thorough assessments of handedness in our future studies. We also plan to explore more precisely the potentials occurring after movement onset. As in many ERP-studies of motor behavior there is no analysis of the components subsequent to the LRP, we recommend paying attention to reafferent and other post-movement MRPs and studying them with regard to their role in feedback processing and sensorimotor integration. We plan to investigate correlations between RAPs and μ-rhythm amplitude. This investigation will shed light on the contribution of movement-related afferent inputs. In this study, we observed irregular weak μ-ERDs in both ME and MI trials, presumably because the trials were short (Pfurtscheller et al., 2008). Moreover, somewhat unexpectedly, we did not find a significant difference in β-ERD during ME versus MI. However, this result could be due to the way the data alignment and averaging was performed: the timing of the ME-related β-ERD is better explained by movement onset, but we had to align the data to stimulus onset because response onset was unknown for MI. This alignment may have resulted in smearing of the averaged ERD time source.

## ACKNOWLEDGEMENTS

This work was supported by the Russian Science Foundation under grant № 21-75-30024.

